# *In silico* analyses identifies sequence contamination thresholds for Nanopore-generated SARS-CoV2 sequences

**DOI:** 10.1101/2023.09.26.559465

**Authors:** Ayooluwa J. Bolaji, Ana T. Duggan

## Abstract

The SARS-CoV-2 pandemic has brought molecular biology and genomic sequencing into the public consciousness and lexicon. With an emphasis on rapid turnaround, genomic data has been used to inform both diagnostic and surveillance decisions for the current pandemic at a previously unheard-of scale. The surge in the submission of genomic data to publicly-available databases has proved essential as comparing different genome sequences offers a wealth of knowledge, including phylogenetic links, modes of transmission, rates of evolution, and the impact of mutations on infection and disease severity. However, the scale of the pandemic has meant that once sequencing runs are performed, they are rarely repeated due to limited sample material and/or the availability of sequencing resources, resulting in some imperfect runs being uploaded to public repositories. As a result, it is crucial to investigate the data obtained from these imperfect runs to determine whether the results are reliable. Numerous studies have identified a variety of sources of contamination in public next-generation sequencing (NGS) data as the number of NGS studies increases along with the diversity of sequencing technologies and procedures [1–3]. For this study, we conducted an *in silico* experiment with known SARS-CoV-2 sequences produced from Oxford Nanopore Technologies sequencing to investigate the effect of contamination on lineage calls and single nucleotide variations (SNVs). Through a series of analyses, we identified a contamination threshold below which runs are expected to generate accurate lineage calls and maintain genomic sequence integrity. Together, these findings provide a benchmark below which imperfect runs may be considered robust for reporting results to both stakeholders and public repositories and reduce the need for repeat or wasted runs.

**Author Summary:** Large-scale genomic comparisons provide a wealth of knowledge, including modes of transmission, rates of evolution, and the impact of mutations on infection, disease severity, and treatment effectiveness. As a result, the public release of genomic data has proven to be crucial. However, studies continue to show that some of the genomic data in public repositories are contaminated due to a variety of reasons. For instance, in the case of SARS-CoV-2 sequences, the pandemic prevented many sequencing runs from being repeated, resulting in some imperfect runs being uploaded to public repositories. It is of note that when genomic data is contaminated, both scientific decisions/studies and public health measures may be compromised. To identify genome contamination threshold(s) for SARS-CoV-2 sequences generated by Nanopore sequencing, computational biology techniques were utilized to generate artificially subsampled contaminated genomes. This is the first study of its kind and so our hope is that the results obtained provide a starting point for the investigation of reporting contamination of NGS data.

## Introduction

Genomics and whole genome sequencing of pathogens provide vital information for disease transmission, identification of novel outbreaks, and vaccine candidate selection [4]. Numerous investigations have shown that in the early days of the COVID-19 pandemic, results from genomic monitoring were not only equivalent to epidemiological contact tracing data, [4] but also capable of tracing previously unidentified linked transmissions [5]. It is noteworthy that public health decisions were guided by genomic investigations in some jurisdictions to stop the spread of SARS-CoV-2, including travel bans and stay-at-home orders[4,6,7]. Thus, it can be concluded that the rapid whole genome sequencing for SARS-CoV-2 is essential for public health intervention.

Since the SARS-CoV outbreak in 2002–2003, genomic information has gained growing importance for addressing outbreaks brought on by pathogenic coronaviruses. Indeed, progress regarding the studies of this virus shifted dramatically as the complete viral genome was sequenced [8]. However, due to the technology available and the lag in data sharing, it took about 3 months to complete the sequencing of the first complete genome of the SARS-CoV virus [9,10]. Complete genomes were generated in 2002-2003 by first propagating the virus in cell lines, extracting viral RNA from these cell lines, and using a Sanger sequencing approach to produce complete and partial genomes [10]. It is worth noting that advances in genomics have significantly improved sequencing methodologies and timelines in less than two decades, owing to the development of third generation NGS and long-read sequencing technologies. Thus, in late December 2019, the first whole genome sequences of the novel beta coronaviruses, now known as SARS-CoV-2, was obtained using metagenomics and NGS approaches - supplemented with PCR and Sanger sequencing [11–13] and made available online within days. The availability of the SARS-CoV-2 reference whole genome sequences facilitated the development of real-time PCR-based diagnostic assays that helped to understand the transmission patterns and epidemiology of the virus [14]. Both partial and whole genome sequences of SARS-CoV-2 genomes have been reported from many parts of the world and these data have proved useful in monitoring the global spread of the virus.

Prior to the 2019-2020 SARS-CoV-2 pandemic, there were approximately 1200 complete betacoronavirus genomes deposited in GenBank. As of July 2023, however, there were over 15.8million sequence submissions of the SARS-CoV-2 genomes available in the Global Initiative on Sharing Avian Influenza Data (GISAID) (https://www.gisaid.org) platform, reflecting a significant increase in the number of available genomes throughout the pandemic. These genomic sequences are generated on different next-generation sequencing (NGS) devices, namely Illumina, Ion Torrent, Oxford Nanopore, and PacBio SMRT platforms. While sequencing technologies have error rates of varying degrees [15,16] genome sequence contamination may also occur during sample preparation and sample processing at both wet and dry lab steps of the workflow. Also, contamination in reference databases is more concerning than contamination in individual sequencing studies and, according to a few studies, human DNA contamination has been found in non-primate reference genomes [2,17]. GenBank has also been reported to contain millions of contaminated sequences, and human contamination in bacterial reference genomes has resulted in thousands of false protein sequences [18]. Therefore, even if researchers properly decontaminated or controlled for contaminants, contamination in reference databases runs the risk of tainting the results of many genomic studies. Further, numerous studies have identified a variety of sources of contamination in public NGS databases and these studies have discovered widespread cross-contamination between samples as well as contamination in sequencing kits and laboratory reagents [18–21].

While NGS has been used for the rapid detection and characterization of positive COVID-19 cases, one of the drawbacks is that NGS runs are rarely repeated for reasons including limited funds to repeat expensive library preparation reactions and NGS remains relatively expensive, even when samples are multiplexed. This has meant that in some cases, the results of some imperfect runs are used to drive public health decisions and are also uploaded to public repositories. Most studies, with few exceptions, do not clearly define the quality control metrics used to include or exclude genomic data from public repositories. Thus, contamination can seriously affect the results of genomic analyses of organisms leading to spurious alignments and incorrect downstream variant calls.

For this study, we conducted an *in silico* experiment using known SARS-CoV-2 sequences produced from Nanopore sequencing. We assessed the effect of contamination on lineage calls and single nucleotide variations (as a measure of genome integrity) using sequences from the same variants and sequences from different variants. The effect of sequencing depth on contamination detection was further investigated using three different numbers of reads (12,500 reads, 25,000 reads, and 50,000 reads) as a measure of sequencing depth. For each sequencing depth, 15 artificially subsampled genomes were generated. These samples were generated by mixing clinical SARS-CoV-2 samples *in silico* at different levels of contamination - low (1% to 9% level) and high (10%, 20%, 30%, 40%, and 50%) contamination levels. Results obtained in this study should help establish internal quality controls and contamination thresholds for SARS-CoV-2 sequences to improve the quality of sequences deposited in public repositories and to offer researchers a standard by which results obtained from contaminated SARS-CoV-2 runs can be trusted for variant calling and other downstream reporting.

## Methods

### SARS-CoV-2 genome sequencing and generation of the i*n silico* contaminated libraries

Amplicons generated using tiling PCR were prepared for Oxford Nanopore Technologies sequencing using the ONT Ligation Sequencing Kit (SQK-LSK109) as per the manufacturer’s guidelines. The resulting reads were basecalled using the Guppy high accuracy model (5.0.7) with default settings. The average number of reads generated for 60 SARS-CoV-2 samples sequenced on a MinION device and 752 samples sequenced on a GridION device were determined using NanoStat (https://github.com/wdecoster/nanostat). The results obtained were used as a guide for the selection of the read lengths as well as experimental design for the generation of the artificial genomes, where low (12,500 reads), medium (25,000 reads), and high (50,000 reads) read depths were explored. Random subsampled artificial sequences were generated with seqtk (https://github.com/lh3/seqtk) for both the background and contaminate samples to represent the artificially contaminated libraries (Table 1). 15 different contamination levels (low levels: 1-9% and high levels: 10%, 20%, 30%, 40%, and 50%) were also studied at each of the three read lengths (Figure 1 and Table 1). It is of note that in this study, the number of reads was used as a measure of sequencing depth.

**Figure 1:**
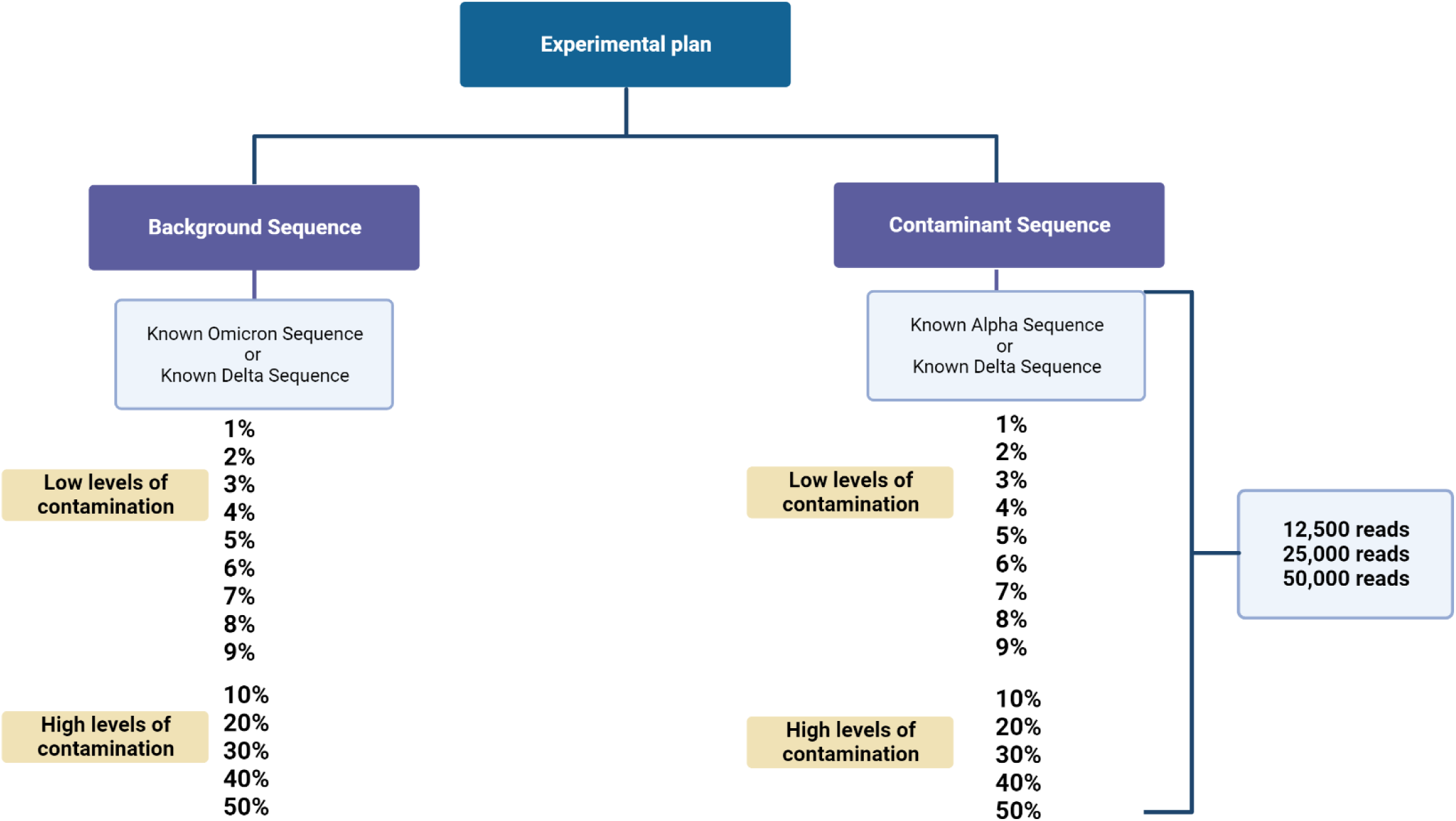
Experimental design of the artificially subsampled genomes for the 15 levels of contamination (low and high levels) at three sequencing depths (low - 12,500 reads, medium - 25,000 reads, and high - 50,000 reads). The controlled datasets were generated from known clinical SARS-CoV-2 samples. Created with BioRender.com.

**Table 1:**
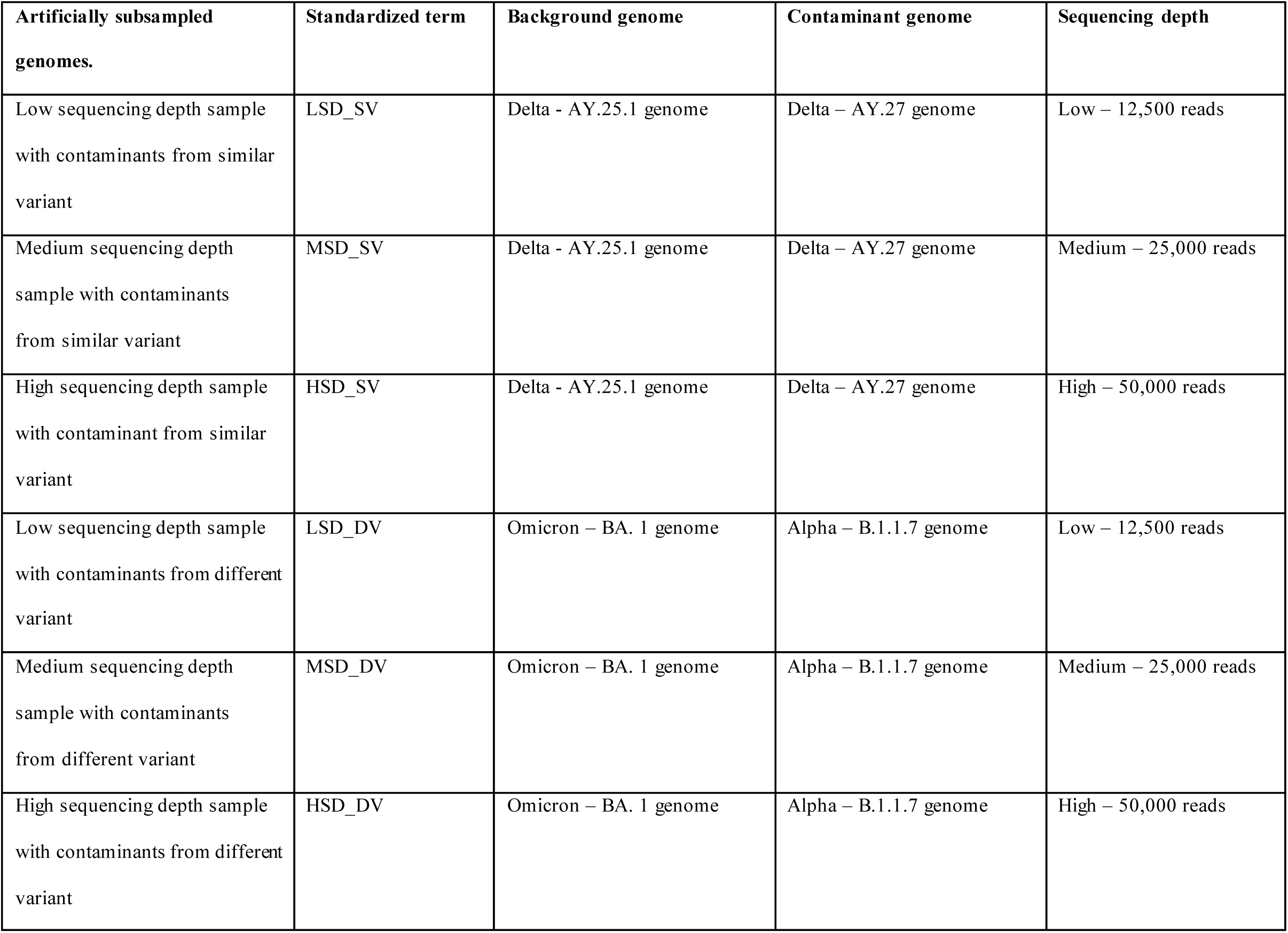
Standardized terms and parameters of the artificially subsampled genomes.

### Data processing

The artificially generated libraries were processed using a nextflow implementation of the ARTIC pipeline (https://github.com/connor-lab/ncov2019-artic-nf). Variant candidates were identified using Nanopolish (https://github.com/jts/nanopolish). Output files generated from the ARTIC pipeline were further processed using ncov-tools to perform quality control on sequencing results (https://github.com/jts/ncov-tools). Reads were mapped to the reference SARS-CoV-2 genome NCBI GenBank accession (MN908947) and lineages were assigned using Pangolin (version 4.0.3, pangoLEARN) (version 1.2.333). The artificially generated datasets (raw reads) as well as their corresponding consensus sequences have been deposited to Zenodo: https://doi.org/10.5281/zenodo.8206455

### Genome pairwise comparison and heat map

Aligned nucleotide consensus genome sequences of both the clinical samples and the artificially generated genomes were imported to MEGA11 software to calculate pairwise distance. The p-distance option was chosen as input for the Model/Method setting while the default options were chosen for the other settings. The pairwise distance output table was imported as a text-delimited file into R v.4.1.1 and the ggplot2 v3.3.1 package was used to generate heat maps for data visualization.

## Results

### Global nucleotide comparison at different levels of contamination for different sequencing depths

To investigate the effect of both low and high levels of contamination on lineage calls and single nucleotide variations as a measure of genome integrity, a series of global nucleotide comparisons using pairwise p-distance analyses were performed. Since the average number of reads for the 768 SARS-CoV-2 clinical samples examined in this study was 46,317 reads and considering the difference in throughput of Nanopore devices (MinION, GridION, and PromethION), three standardized read lengths or run depths were chosen as a measure of sequencing depth– low (12,500 reads), medium (25,000 reads), and high (50,000 reads). Samples were generated through *in silico* artificial mixtures of reads to simulate contaminated libraries of controlled datasets generated from clinical samples. The distance (proportion) of nucleotide sites was compared and plotted as a heat map for all artificially generated samples at the three sequencing depths – low (12,500 reads), medium (25,000 reads), and high (50,000 reads) (Figure 2). This comparison was done for samples contaminated by reads generated from both similar (Figure 2A) and different SARS-CoV-2 viral strains (Figure 2B). The results obtained show that for global nucleotide comparison, regardless of the sequencing depth and the contamination types (i.e., similar (Figure 2A) or different variant contaminants (Figure 2B)), differences observed for global nucleotide composition among the samples were not substantial for contamination levels less than 20% (see Figure 2 for the low sequencing depth, supplementary Figures 1 and 2 for medium and high sequencing depths).

**Figure 2.**
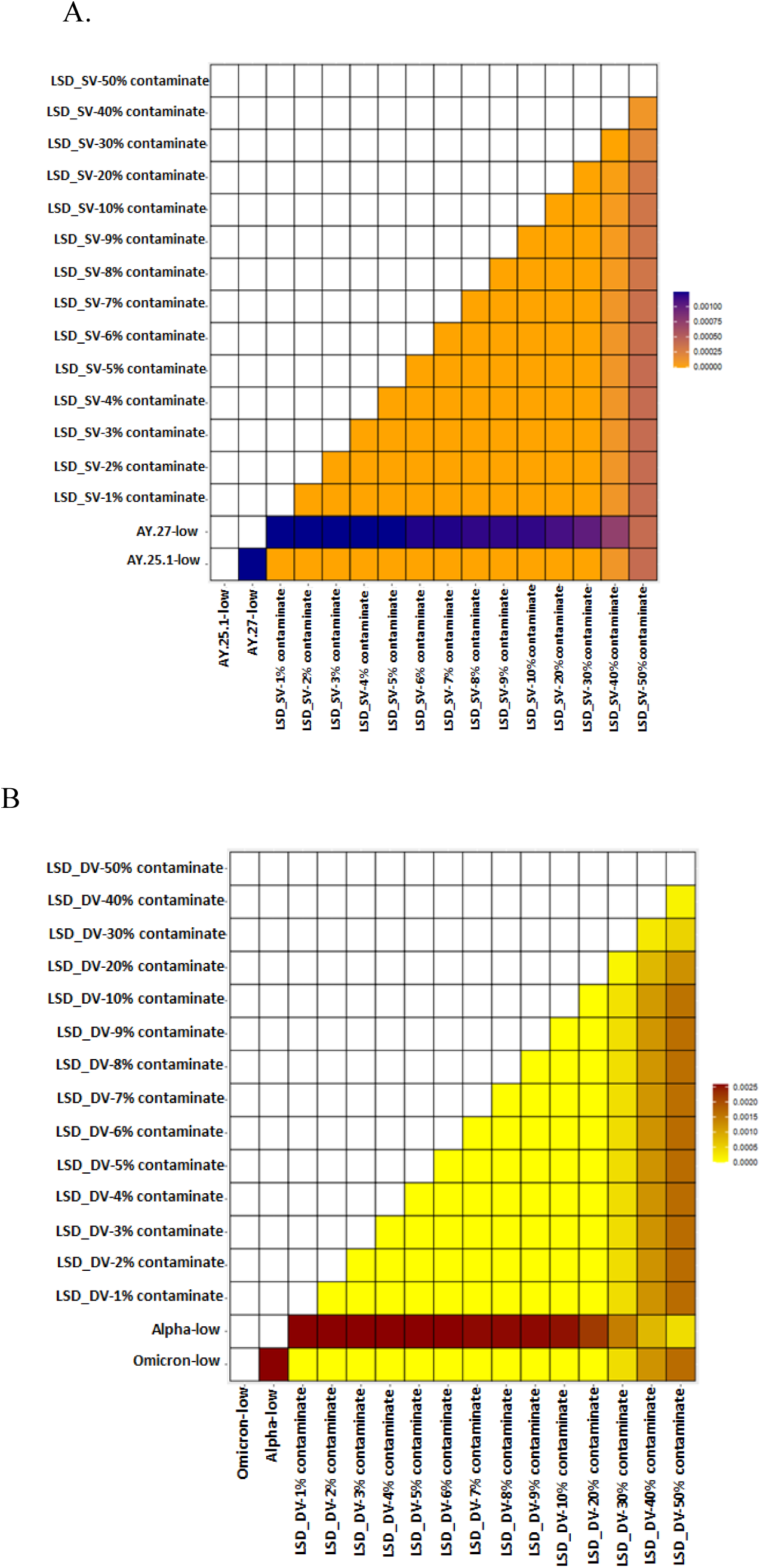
Global nucleotide comparison of artificially generated contaminated samples and their corresponding background clinical samples at a low sequencing depth. A) A heatmap of the pairwise p-distance comparison of the LSD samples - a delta background sequence (AY.25.1) contaminated with a similar delta contaminant sequence (AY.27). B) A heatmap of the pairwise p-distance comparison of the HSD samples – an omicron background sequence (BA.1) contaminated with an alpha contaminant sequence (B.1.1.7).

### The effect of contamination from similar variants on genome integrity and lineage calls

The impacts of contamination on single nucleotide variations (SNVs) and lineage call outputs for the SARS-CoV-2 genome were assessed by creating *in silico* artificial mixtures of reads to simulate contaminated genomes. By subsampling the sequences of a known clinical delta sample (AY.25.1) contaminated with reads from another known clinical delta sample (AY.27), 15 different contamination scenarios were simulated to quantify the effect of contamination. Phylogenetic trees were constructed to examine the impact of single nucleotide variations (SNVs) found within each subsampled dataset and sequences from the clinical samples served as controls. The identified SNVs were plotted with an associated single nucleotide polymorphism (SNP) matrix (Figure 3 and Supplementary Figures 3 & 4). Seven quality control metrics (QC metrics) were highlighted as important metrics in determining contamination thresholds and the effect(s) of sequence contamination on genome completeness and integrity. These metrics include the number of consensus single nucleotide variations (SNVs), the number of consensus ‘n’, the number of variants SNVs, the number of variants indel, genome completeness, lineage, and Scorpio call.

**Figure 3.**
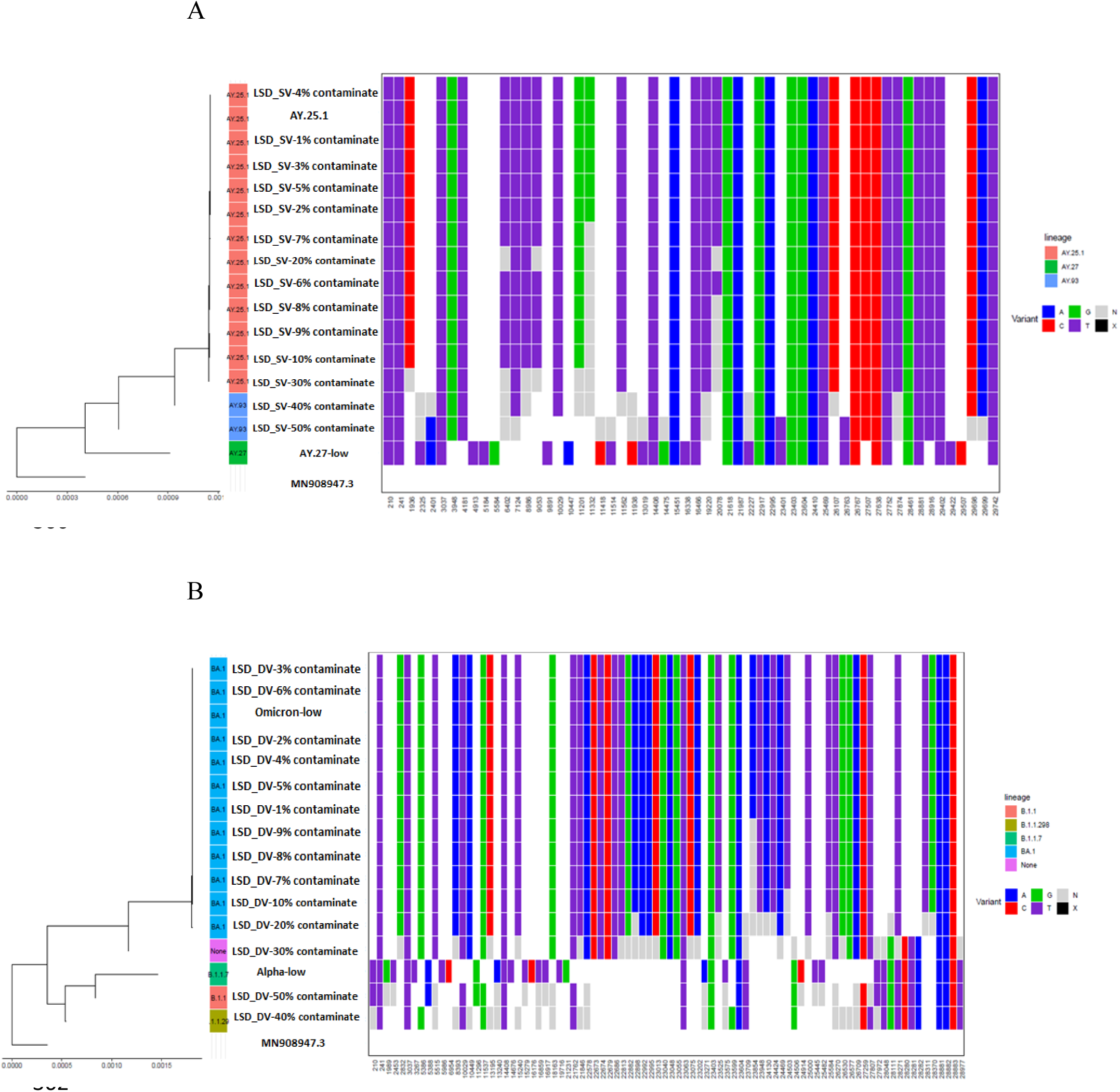
Phylogenetic tree and heatmap showing single nucleotide variations (SNVs) at different positions of the SARS-CoV-2 genome for (A) AY.25.1 (delta variant) contaminated with an AY.27 (delta variant) sequence at contamination levels 1 -10%, 20%, 30%, 40%, and 50%. (B) BA.1 (omicron variant) low sequencing depth sequence (12,500 reads) contaminated with a B.1.1.29 (alpha variant) sequence at contamination levels 1 -10%, 20%, 30%, 40%, and 50%.

We examined the 15 artificial samples generated from an AY.25.1 clinical delta sample (background sequence) and an AY.27 clinical delta sample (contaminate sample) for all samples, at a low sequencing depth (12,500 reads). Changes in both the numbers of consensus SNVs and consensus ‘N’ (number of missing data) were investigated as these two metrics are essential determinants of genome integrity and completeness. For the LSD_SV genomes (12,500 reads), differences in the two aforementioned metrics were observed for the genomes with contamination levels greater than 5% (625 reads) (Figure 3A, Table 2) – wherein as the levels of contamination increased, a decrease in the number of SNVs and an increase in the number of consensus ‘N’s compared to the clinical control samples (Figure 3A, Table 2). Further, it was observed that the LSD_SV genomes were assigned incorrect lineage calls at contamination levels greater than 30% (3,750 reads). Thus, for LSD_SV genomes, the contamination threshold for preserving genome integrity is 5% while the identified threshold for lineage calls is 30% (Figure 3A, Table 2). For the MSD_SV genomes a decrease in the number of SNVs (from 40 to 39) and an increase in the number of consensus ‘n’ (from 189 to 190) were observed at contamination levels greater than 4% (1,000 reads) while for lineage calls, the identified threshold for contamination was 30% (7,500 reads) (Supplementary Figure 3A and Supplementary Table 1A). Lastly, For the HSD_SV samples (50,000 reads), a contamination threshold of 10% (5,000 reads) was identified for SNVs and a threshold of 50% (25,000 reads), was identified for lineage calls (Supplementary Figure 3B and Supplementary Table 1B). In conclusion, for contamination by a similar SARS-CoV-2 variant, the contamination threshold identified for lineage call was 30% for both LSD_SV and MSD_SV and 50% for HSD_SV genomes. However, for genome integrity, the contamination threshold was 5% for low, 4% for medium, and 10 % for high sequencing depths (Figures 4 A & B).

**Figure 4.**
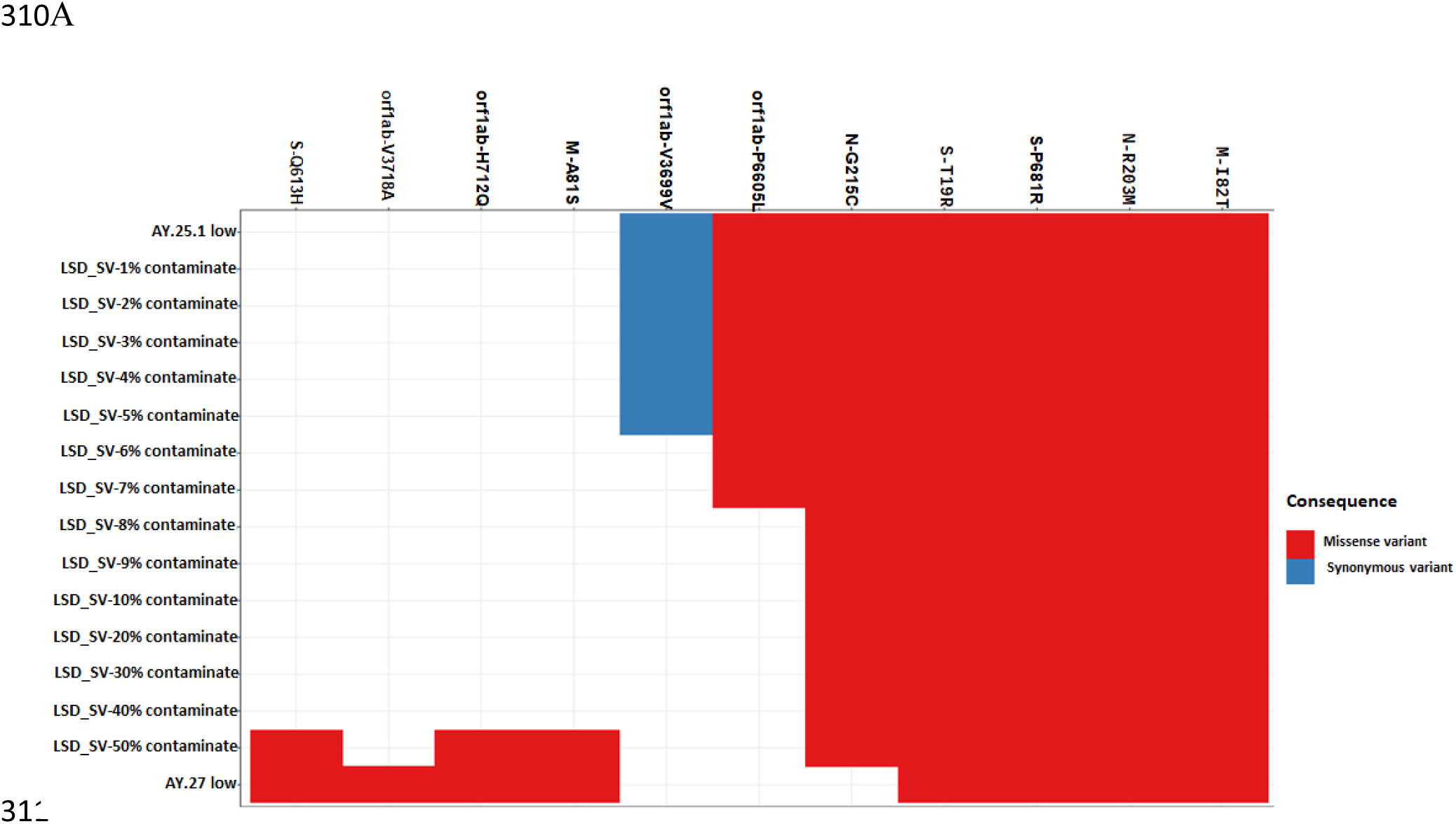
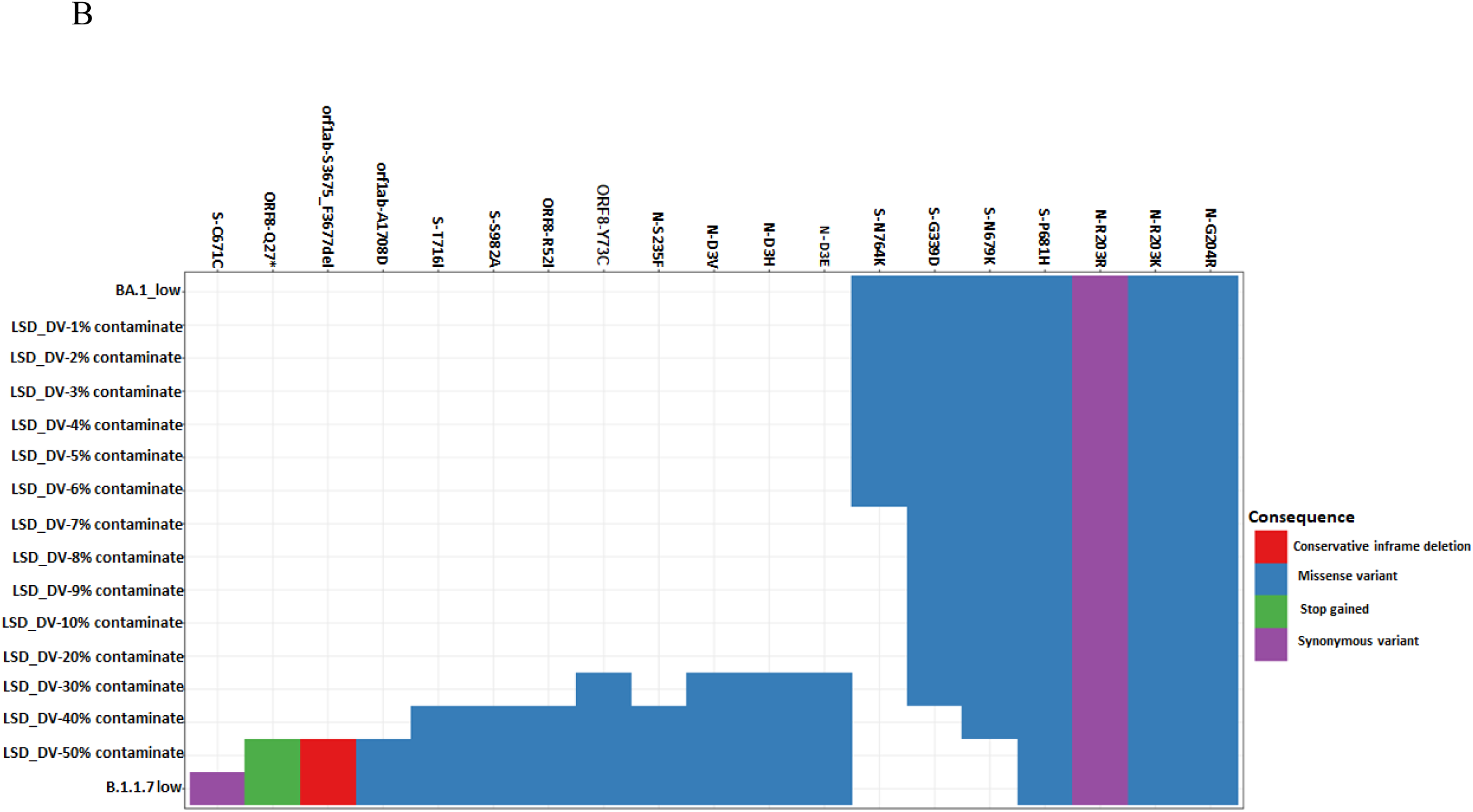
Mutational profile comparison of SARS-CoV-2 genome for the clinical genomes to the artificially generated genomes for (A) LSD_SV (AY.25.1 contaminated with an AY.27 variant) sequence at contamination levels 1-10%, 20%, 30%, 40%, and 50%. (B) LSD_DV (BA.1 contaminated with a B.1.1.29 variant at contamination levels 1-10%, 20%, 30%, 40%, and 50%.

**Table 2:** Quality control metrics comparison for artificially subsampled genomes of contamination by similar variants at a low sequencing depth – all LSD_SV genomes.

| Genome | Num of<br>consensus<br>snvs | Number of<br>consensus 'n' | Number of<br>variants<br>SNVs | Number of<br>variants indel | Number of<br>variants<br>indel<br>triplet | Mean<br>sequencing<br>depth | Median<br>sequence<br>depth | Scaled<br>variants<br>SNVs | Genome<br>completeness | Linea<br>ge | Lineage<br>note | Scorpio<br>calls | Watch<br>mutations |
| --- | --- | --- | --- | --- | --- | --- | --- | --- | --- | --- | --- | --- | --- |
| AY.25.1_low | 40 | 190 | 45 | 2 | 2 | 472.1 | 452 | 45.29 | 0.9936 | AY.25<br>.1 | alt/ref/am<br>b:13/0/0 | Delta<br>(B.1.617.<br>2-like) | S:G142D,S:L4<br>52R |
| LSD_S -1 %<br>contaminate | 40 | 190 | 45 | 2 | 2 | 472.1 | 454 | 45.29 | 0.9936 | AY.25<br>.1 | alt/ref/am<br>b:13/0/0 | Delta<br>(B.1.617.<br>2-like | S:G142D,S:L4<br>52R |
| LSD_SV-2%<br>contaminate | 40 | 190 | 45 | 2 | 2 | 472.1 | 448 | 45.29 | 0.9936 | AY.25<br>.1 | alt/ref/am<br>b:13/0/0 | Delta<br>(B.1.617.<br>2-like | S:G142D,S:L4<br>52R |
| LSD_SV-3%<br>contaminate | 40 | 190 | 45 | 2 | 2 | 472.1 | 450 | 45.29 | 0.9936 | AY.25<br>.1 | alt/ref/am<br>b:13/0/0 | Delta<br>(B.1.617.<br>2-like | S:G142D,S:L4<br>52R |
| LSD_SV-4%<br>contaminate | 40 | 190 | 45 | 2 | 2 | 472.1 | 446 | 45.29 | 0.9936 | AY.25<br>.1 | alt/ref/am<br>b:13/0/0 | Delta<br>(B.1.617.<br>2-like | S:G142D,S:L4<br>52R |
| LSD_SV-5%<br>contaminate | 40 | 190 | 45 | 2 | 2 | 472.1 | 448 | 45.29 | 0.9936 | AY.25<br>.1 | alt/ref/am<br>b:13/0/0 | Delta<br>(B.1.617.<br>2-like | S:G142D,S:L4<br>52R |
| LSD_SV-6%<br>contaminate | 39 | 190 | 44 | 3 | 2 | 472.1 | 442 | 44.28 | 0.9936 | AY.25<br>.1 | alt/ref/am<br>b:13/0/0 | Delta<br>(B.1.617.<br>2-like | S:G142D,S:L4<br>52R |
| LSD_SV-7%<br>contaminate | 39 | 190 | 44 | 3 | 2 | 472.1 | 441 | 44.28 | 0.9936 | AY.25<br>.1 | alt/ref/am<br>b:13/0/0 | Delta<br>(B.1.617.<br>2-like | S:G142D,S:L4<br>52R |
| LSD_SV-8%<br>contaminate | 38 | 191 | 43 | 3 | 2 | 472.1 | 444 | 43.28 | 0.9936 | AY.25<br>.1 | alt/ref/am<br>b:13/0/0 | Delta<br>(B.1.617.<br>2-like | S:G142D,S:L4<br>52R |
| LSD_SV-9%<br>contaminate | 38 | 191 | 43 | 3 | 2 | 472.2 | 445 | 43.28 | 0.9935 | AY.25<br>.1 | alt/ref/am<br>b:13/0/0 | Delta<br>(B.1.617.<br>2-like | S:G142D,S:L4<br>52R |
| LSD_SV-10%<br>contaminate | 38 | 191 | 43 | 3 | 2 | 472.1 | 449 | 43.28 | 0.9934 | AY.25<br>.1 | alt/ref/am<br>b:13/0/0 | Delta<br>(B.1.617.<br>2-like | S:G142D,S:L4<br>52R |
| LSD_SV-20%<br>contaminate | 36 | 194 | 41 | 2 | 2 | 472.2 | 444 | 41.27 | 0.9935 | AY.25<br>.1 | alt/ref/am<br>b:13/0/0 | Delta<br>(B.1.617.<br>2-like | S:G142D,S:L4<br>52R |
| LSD_SV-30%<br>contaminate | 33 | 196 | 38 | 3 | 2 | 472.4 | 438 | 38.25 | 0.9934 | AY.25<br>.1 | alt/ref/am<br>b:13/0/0 | Delta<br>(B.1.617.<br>2-like | S:G142D,S:L4<br>52R |
| LSD_SV-40%<br>contaminate | 29 | 202 | 34 | 2 | 2 | 472.3 | 433 | 34.23 | 0.9932 | AY.93 | alt/ref/am<br>b:13/0/0 | Delta<br>(B.1.617.<br>2-like | S:G142D,S:L4<br>52R |
| LSD_SV-50%<br>contaminate | 28 | 203 | 32 | 2 | 2 | 472 | 439 | 32.22 | 0.9932 | AY.93 | alt/ref/am<br>b:13/0/0 | Delta<br>(B.1.617.<br>2-like | S:G142D,S:L4<br>52R |
| AY.27_low | 39 | 189 | 43 | 3 | 2 | 471.7 | 454 | 43.27 | 0.9937 | AY.27 | alt/ref/am<br>b:13/0/0 | Delta<br>(B.1.617.<br>2-like | S:G142D,S:L4<br>52R |

### The effect of contamination on genome integrity and lineage calls for SARS-CoV-2 sequences for different variants

The 15 *in silico* samples were also generated by artificially subsampling sequences from an omicron clinical sample (BA.1), contaminated with an alpha clinical sample (B.1.1.7). We investigated the effect of different levels of contamination on SARS-CoV-2 sequences contaminated by different strains. A contamination threshold was identified for changes in SNVs and the number of consensus ‘N’-a measure of genome integrity and lineage calls. Three sequencing depths – low (12,500 reads), medium (25,000 reads), and high (50,000 reads) were examined.

It was observed that at a low sequencing depth (12,500 reads), the number of consensus SNVs for the clinical omicron BA.1 sample was 56, the number of consensus ‘ N’ as a measure of missing nucleotide was 189 and the number of variants SNVs was 61 (Table 3). Therefore, differences in these QC metrics were investigated for each of the artificially generated genomes. For the LSD_DV at a contamination level of 7%, it was observed that the number of consensus SNVs changed from 56 to 55, the number of consensus ‘N’ changed to 190 while the number of variant SNVs remained at 60 and other QC metrics remained unchanged at this contamination level (Table 3). However, at 30%, the assigned lineage calls for the artificially generated genome (LSD_DV) changed from BA.1 to none (Table 3), and this held true for artificial genomes with 40% and 50% contamination. Taken together, for low sequencing depth (LSD_DV), 6% level of contamination (750 reads) was identified as the contamination threshold for the preservation of genome integrity while a 20% level of contamination (2,500 reads) was identified as the threshold for accurate lineage call (Figure 3B, Table 3). For the MSD_DV samples (25,000 reads), a decrease in the number of consensus SNVs (from 56 to 55), an increase in the number of consensus ‘N’ (189 to 190), and a decrease in the number of variant SNVs (61 to 60) were observed as the contaminant levels increased above 7% (Supplementary Figure 4A and Supplementary Table 2A). Also, at a 30% level of contamination for MSD_DV samples, the assigned lineage calls for samples changed from BA.1 to unassigned, and this was equally observed for samples with both 40% and 50% levels of contaminants. Therefore, at a medium sequencing depth (25,000 reads), the contamination threshold for preserving genome integrity was identified to be 7% while the contamination threshold for lineage call was 20%. For the HSD_DV samples (50,000 reads), the artificially generated genome with an 8% and above level of contamination, showed a decrease in the number of consensus SNVs (from 55 to 54), an increase in the number of consensus ‘N’ (from 189 to 190), and a decrease in the number of variants SNVs (from 61 to 60) (Supplementary Figure 4B). Also, changes in lineage call assignment were not observed until the contamination threshold reached 30% (lineage call assignment changed from BA.1 to an unassigned lineage) (Supplementary Table 2B). In conclusion, for artificial genomes generated by mixing different SARS-CoV-2 variants (an omicron sample contaminated by an alpha sample), the contamination threshold identified for lineage call was 20% at all sequencing depths while for genome integrity, the contamination threshold identified for LSD (12,500 reads) was 6% and 7% for both MSD (25,000 reads) and HSD (50,000 reads) depths (Figures 5 A & B).

**Table 3:** Quality control metrics comparison for artificially subsampled genomes of contamination by different variants at a low sequencing depth – for all LSD_DV genomes.

| Genome | Num. of<br>consensus<br>_snvs | Number of<br>consensus ‘n’ | Number of<br>variants<br>SNVs | Number of<br>variants<br>indel | Number of<br>variants<br>indel triplet | Mean<br>sequencing<br>depth | Median<br>sequence<br>depth | Scaled<br>variants<br>SNVs | Genome<br>completeness | Lineage | Lineage<br>note | Scorpio<br>calls | Watch mutations |
| --- | --- | --- | --- | --- | --- | --- | --- | --- | --- | --- | --- | --- | --- |
| BA.1_low | 56 | 189 | 61 | 7 | 7 | 470.4 | 434 | 61.39 | 0.9937 | BA.1 | alt/ref/a<br>mb:55/0/<br>3 | Omicron<br>(BA.1-like) | S:del69-70,S:K417N,S:Q4<br>93R,S:N501Y,S:P<br>681H,S:P681H11 |
| LSD_DV-1%<br>contaminate | 56 | 189 | 61 | 7 | 7 | 470.4 | 440 | 61.39 | 0.9937 | BA.1 | alt/ref/a<br>mb:55/0/<br>3 | Omicron<br>(BA.1-like) | S:del69-70,S:K417N,S:Q4<br>93R,S:N501Y,S:P<br>681H,S:P681H |
| LSD_DV-2%<br>contaminate | 56 | 189 | 61 | 7 | 7 | 470.4 | 442 | 61.39 | 0.9937 | BA.1 | alt/ref/a<br>mb:55/0/<br>3 | Omicron<br>(BA.1-like) | S:del69-70,S:K417N,S:Q4<br>93R,S:N501Y,S:P<br>681H,S:P681H |
| LSD_DV-3%<br>contaminate | 56 | 189 | 61 | 7 | 7 | 470.4 | 452 | 61.39 | 0.9937 | BA.1 | alt/ref/a<br>mb:55/0/<br>3 | Omicron<br>(BA.1-like) | S:del69-70,S:K417N,S:Q4<br>93R,S:N501Y,S:P<br>681H,S:P681H |
| LSD_DV-4%<br>contaminate | 56 | 189 | 61 | 7 | 7 | 470.4 | 455 | 61.39 | 0.9937 | BA.1 | alt/ref/a<br>mb:55/0/<br>3 | Omicron<br>(BA.1-like) | S:del69-70,S:K417N,S:Q4<br>93R,S:N501Y,S:P<br>681H,S:P681H |
| LSD_DV-5%<br>contaminate | 56 | 189 | 61 | 7 | 7 | 470.4 | 449 | 61.39 | 0.9937 | BA.1 | alt/ref/a<br>mb:55/0/<br>3 | Omicron<br>(BA.1-like) | S:del69-70,S:K417N,S:Q4<br>93R,S:N501Y,S:P<br>681H,S:P681H |
| LSD_DV-6%<br>contaminate | 56 | 189 | 61 | 7 | 7 | 470.4 | 454 | 61.39 | 0.9937 | BA.1 | alt/ref/a<br>mb:55/0/<br>3 | Omicron<br>(BA.1-like) | S:del69-70,S:K417N,S:Q4<br>93R,S:N501Y,S:P<br>681H,S:P681H |
| LSD_DV-7%<br>contaminate | 55 | 190 | 60 | 7 | 7 | 470.4 | 454 | 60.39 | 0.9936 | BA.1 | alt/ref/a<br>mb:54/0/<br>3 | Omicron<br>(BA.1-like) | S:del69-70,S:K417N,S:Q4<br>93R,S:N501Y,S:P<br>681H,S:P681H |
| LSD_DV-8%<br>contaminate | 55 | 190 | 60 | 7 | 7 | 470.4 | 455 | 60.39 | 0.9936 | BA.1 | alt/ref/a<br>mb:54/0/<br>3 | Omicron<br>(BA.1-like) | S:del69-70,S:K417N,S:Q4<br>93R,S:N501Y,S:P<br>681H,S:P681H |
| LSD_DV-9%<br>contaminate | 55 | 190 | 60 | 7 | 7 | 470.4 | 456 | 60.39 | 0.9935 | BA.1 | alt/ref/a<br>mb:54/0/<br>3 | Omicron<br>(BA.1-like) | S:del69-70,S:K417N,S:Q4<br>93R,S:N501Y,S:P<br>681H,S:P681H |
| LSD_DV-10%<br>contaminate | 54 | 191 | 59 | 7 | 7 | 470.4 | 465 | 59.38 | 0.9936 | BA.1 | alt/ref/a<br>mb:53/0/<br>3 | Omicron<br>(BA.1-like) | S:del69-70,S:K417N,S:Q4<br>93R,S:N501Y,S:P<br>681H,S:P681H |
| LSD_DV-20%<br>contaminate | 46 | 210 | 51 | 6 | 6 | 470.6 | 452 | 51.36 | 0.993 | BA.1 | alt/ref/a<br>mb:46/0/<br>3 | Omicron<br>(BA.1-like) | S:del69-70,S:K417N,S:Q4<br>93R,S:N501Y,S:P<br>681H,S:P681H |
| LSD_DV-30%<br>contaminate | 34 | 214 | 38 | 5 | 5 | 470.7 | 453 | 38.28 | 0.9928 | None | alt/ref/a<br>mb:20/5/<br>BA.1.13 | Probable<br>Omicron<br>(Unassigned<br>) | S:del69-70,S:Q493R,S:N5<br>01Y,S:P681H,S:P<br>681H |
| LSD_DV-40%<br>contaminate | 25 | 214 | 28 | 4 | 3 | 470.7 | 464 | 28.2 | 0.9928 | B.1.1.298 | none |  | S:del69-70,S:N501Y,S:P6<br>81H,S:P681H,S:T<br>716I,S:S982A |
| LSD_DV-50%<br>contaminate | 26 | 210 | 28 | 3 | 2 | 471 | 451 | 28.2 | 0.993 | B.1.1 | none |  | S:del69-70,S:N501Y,S:P6<br>81H,S:P681H,S:T<br>716I,S:S982A |
| **B.1.1.7 low** | 40 | 192 | 44 | 4 | 3 | 471.1 | 476 | 44.28 | 0.9936 | B.1.1.7 | alt/ref/a<br>mb:21/1/<br>0 | Alpha<br>(B.1.1.7-like) | S:del69-70,S:del144,S:N5<br>01Y,S:A570D,S:P<br>681H,S:P681H,S:<br>T716I,S:S982A,S:<br>D1118H |

**Table 4:** A summary of the identified threshold for the artificially subsampled genomes as well as the origin of both background and contaminant samples.

| Standardized term | The identified threshold for genome integrity call | The identified threshold for lineage call |
| --- | --- | --- |
| LSD_SV | 5 percent | 30 percent |
| MSD_SV | 4 percent | 30 percent |
| HSD_SV | 10 percent | 50 percent |
| LSD_DV | 6 percent | 20 percent |
| MSD_DV | 7 percent | 20 percent |
| HSD_DV | 7 percent | 20 percent |

**Figure 5.**
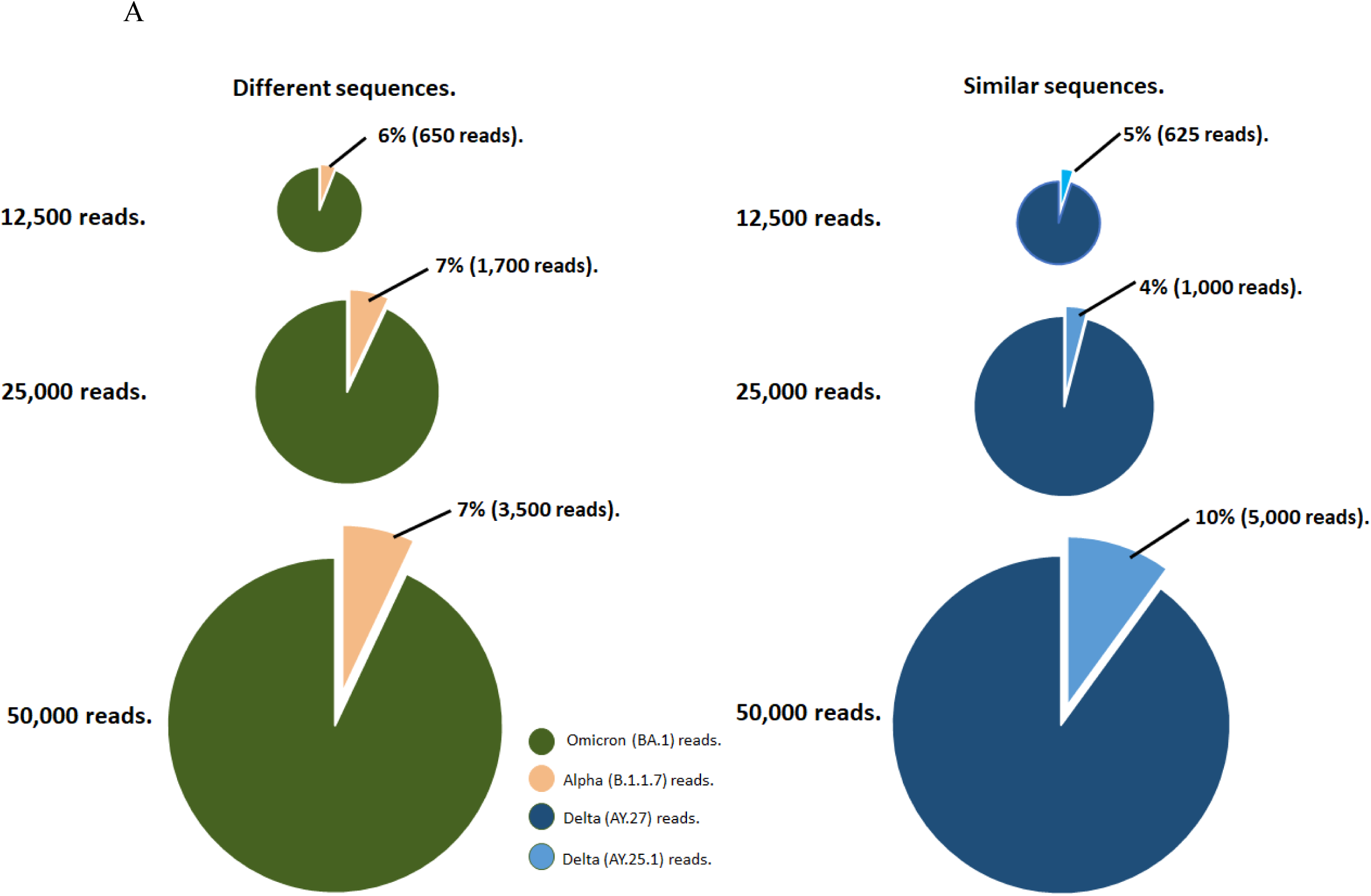
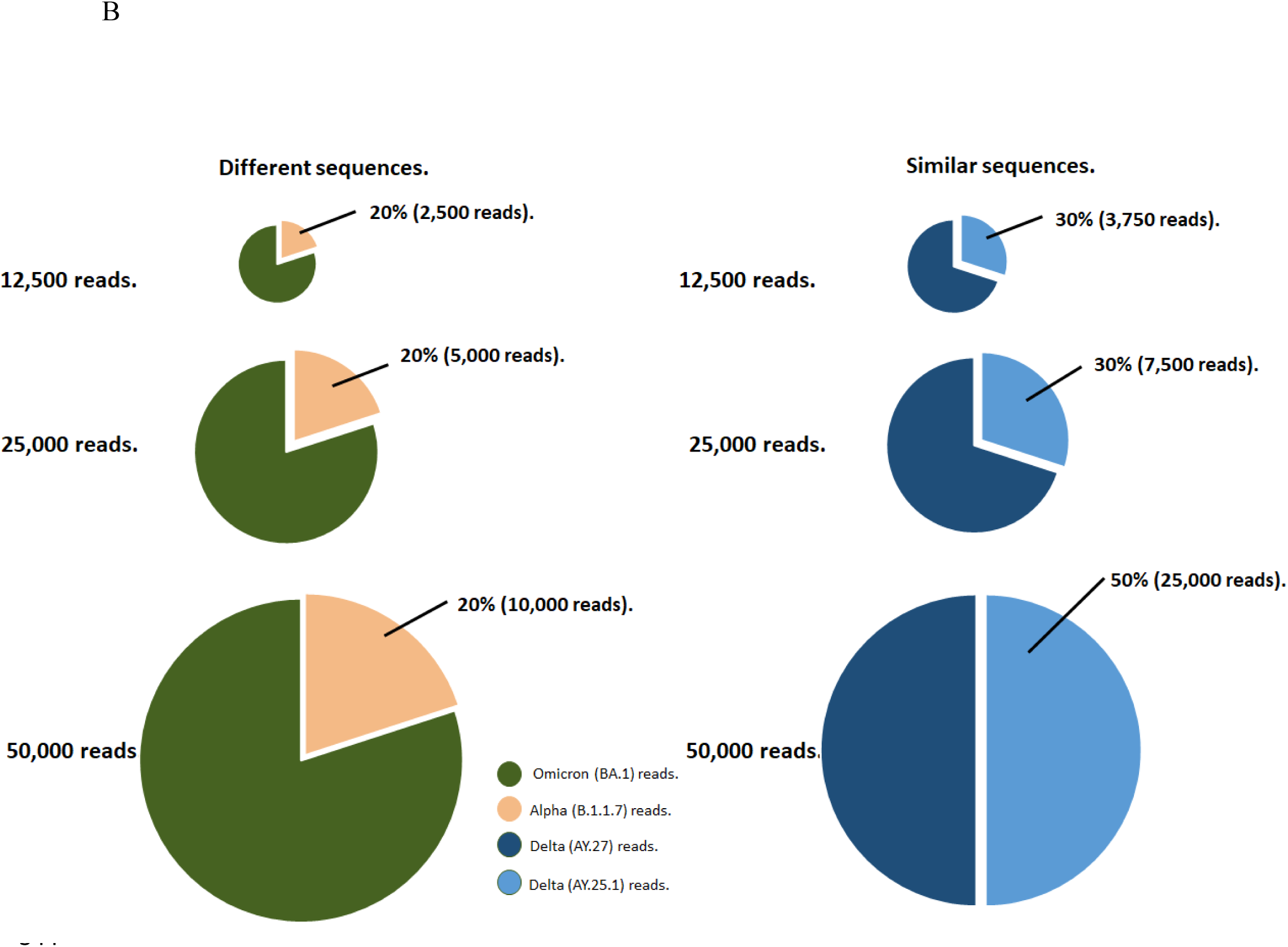
A model for sequence contamination threshold and summary of the findings of this study, depicting: A) the contamination threshold for maintaining genome integrity at low (12,500 reads), medium (25,000 reads), and high (50,000 reads) sequencing depths for both similar and different sequences. B) the contamination threshold identified to maintain an accurate lineage call at low (12,500 reads), medium (25,000 reads), and high (50,000 reads) sequencing depths for different sequences.

## Discussion

The scale of sequencing data available in public repositories over the course of the SARS-CoV-2 pandemic is unprecedented. Due to the rapidly evolving nature of the SARS-CoV-2 genome, routine monitoring and public health warnings were crucial in controlling the pandemic. Continuous monitoring and genomic sequencing during the SARS-CoV-2 coronavirus pandemic also hastened the development of the most effective vaccines [22]. However, recurrent mutations in the SARS-CoV-2 genome have tested the efficacy of the vaccines and point to the need for routine updates to both the vaccine targets and vaccination schedules [22,23]. The importance of routine monitoring of SARS-CoV-2 mutations for public health applications cannot be overstated, therefore it is critical that we maintain confidence in the sequences both submitted and pulled from public repositories lest erroneous variants aff ect major public health decisions [24].

Contaminant-induced mutations have been found and documented in other large-scale genomic studies and it was concluded that these contaminated sequences can spread into and across databases over time [2]. This issue cannot be ignored since genome sequences are frequently obtained from these public repositories/databases, based on the types of sequences they contained. Therefore, researchers interested in a particular genome can collect hundreds of sequences for comparative genomic or phylogenomic investigations in this manner. Lupo et al. demonstrated the presence of mis-affiliated genomes in NCBI RefSeq [1]. While these genomes may not be contaminated in the strictest sense, the dominant organism was not what was expected in the study, leading to problems for downstream analyses and reporting [1]. Despite the findings of these studies, sequences submitted to public repositories/databases are rarely checked for contamination [1]. To further validate the effect of contamination on sequencing data and demonstrate the need for contaminant investigation before data are uploaded to p ublic repositories, this study aimed to identify a contamination threshold for which runs can be considered ideal for upload to public repositories while also offering practical guidelines.

Since there is no consensus within the scientific community on how to validate genome integrity, we investigated the amino acid mutational profile, genome completeness, number of SNVs, number of consensus n, number of variant SNVs, and indels for all samples as a measure of genome integrity for this study (Tables 1&2; Supplementary Tables 1&2). Further, to identify the differences between the clinical samples and the artificially subsampled genomes, we generated an amino acid mutation heatmap. As mutational profiles and other host-modulating factors have been reported as major contributors to disease severity in COVID-19 [25], there is a critical need to evaluate the effect of contamination on mutational profiles that may be of clinical importance. The mutational profile compared all defining mutations of the artificially subsampled genomes to the clinical samples and also identified the type/nature of the mutations (conservative in-frame deletion, disruptive in-frame deletion, missense variant, stop gained, and synonymous variant) (Figure 4). We believe that by examining the mutational profile of the samples, the similarities, and differences present in each sample, compared to each other, may be determined. The amino acid mutation plots reveal the similarity in the mutation profile of each artificially subsampled and the clinical control samples. Samples that have similar genome composition (LSD_SV, MSD_SV, and HSD_SV) also had similar mutational profiles while samples with contaminants from different variants (LSD_DV, MSD_DV, and HSD_DV) had different mutational profiles (Table 1 and 4). It is noteworthy that the artificially subsampled genomes that contained less than 6% of contaminant from a substrain of the same variant had similar mutational profiles to the clinical samples at all levels of contaminations. While the artificially generated subsampled genomes that contained less than 7% of contaminant from a divergent variant had similar mutational profiles to the corresponding clinical samples at all levels of contamination.

We further investigated the effect of contamination on the phylogenetic placement and sample relatedness of the artificially subsampled genomes (Figure 2; Supplementary Figures 3&4). The results obtained from the phylogenetic analyses are in agreement with the ide ntified contamination thresholds for mutation profile as a measure of genome integrity, wherein the artificially subsampled genomes with contaminants of less than 5% for LSD_SV and 6% for LSD_DV clustered in the same branch with the corresponding clinical samples. Similar results were also obtained for both MSD_SV and HSD_SV as well as for MSD_DV and HSD_DV. With this observation, we showed that at contamination levels of less than 6%, at all sequencing depths, the artificially subsampled genomes were closely related to the clinical samples. Thus, we concluded that contamination levels of 5% and below do not affect genome-relatedness and integrity.

By performing a global nucleotide comparison, varying both the levels of simulated contamination and the sequencing depth, we investigated the effect of contamination on the artificially subsampled genomes. According to the results obtained from the p-distance pairwise comparison analysis, irrespective of the sequencing depth and the contamination types (i.e., contaminants from a substrain of the same variant or a different variant), differences observed for global nucleotide composition among the samples were not substantial for contamination levels less than 20% when the metric of interest is simply the lineage assignment. Since p-distance is the proportion of nucleotide sites at which two sequences being compared are different, this result is expected. The analysis performed considers all nucleotides present in the samples compared without any regard for the origin of the nucleotide (i.e., contaminant or not). However, it is noteworthy that with contamination levels greater than 20%, differences were observed at the global nucleotide levels when compared to the original clinical samples at all sequencing depths for both types of contaminants (Figure 2 and Supplementary Figures 1&2).

Some studies have identified the importance of lineage tracking and its role in providing answers to evolutionary questions about the SARS-CoV-2 genome [26,27]. The extensive recombination between SARS-CoV-2 strains, first identified by so-called “deltacron” lineages with diagnostic mutations associated with both the delta and omicron variants have become identified with increasing frequency since late 2021, and the emergence of the omicron variant [28,29]. Thus, the accurate assignment of lineage calls for SARS-CoV-2 lineages is important, coupled with the fact that these lineages also offer insights for clinicians and pub lic health personnel during an outbreak of infection. Based on the above notion, we investigated the effect that the different levels and types of contaminations had on the accuracy of lineage calls (Tables 1&2; Supplementary Tables 1&2). Our results showed that regardless of the type of contaminant (similar or different sequences), a 20% contamination threshold was the maximum amount of contaminant permissible for accurate linage calls (Tables 1&2; Supplementary Tables 1&2).

It has been observed that foreign sequences can be introduced at many different stages of the sequencing process, from organism culture to data processing [2]. Here, we offer some practical guidelines on how to track contaminants during sequencing experiments. We recommend that researchers include a negative control in the following steps: (i) nucleic acid extraction, (ii) nucleic acid amplification (if applicable), and (iii) library preparation steps. By having multiple negative controls introduced at different stages of the sequencing experiment, the source of contamination may be identified. It is also recommended that these negative controls be carried forward to the data processing steps so that if contamination occurs, the amount of sequenced data present in the negative controls could be investigated and used to determine the appropriate contamination threshold based on the objective(s) of the sequencing experiment in question.

In conclusion, given that this study is the first of its kind, we are aware that these identified thresholds may change as more sequence data become available and as more studies expand on and investigate the parameters required for genome integrity and lineage calls. However, we hope that having a standardized method for determining the integrity of genomes and lineage calls will provide a benchmark below which imperfect runs may be considered robust for reporting results to both stakeholders and public repositories thereby reducing the need for repeat or wasted runs. In this study, we investigated contamination thresholds for SARS-CoV-2 samples generated by Nanopore sequencing by conducting *in silico* analyses. A contamination threshold of 5% was identified wherein the integrity of the genome was not compromised and a contamination threshold of 20% for lineage calls. Our results suggest that a stricter threshold should be established if the preservation of genome integrity is of utmost importance. Future larger-scale studies are warranted to systematically investigate the effects of contamination on both SARS-CoV-2 reads and other viral and bacterial sequences to serve as a check step for sequencing upload.

## Acknowledgements

We owe a debt of gratitude to Dr. Anna Majer and the DNA core team at the National Microbiology Laboratory for sequencing the clinical samples utilized in this study. We sincerely thank Dr. Andrea Tyler for her contributions to the experimental design of this study and we appreciate the insightful discussions had with Dr. David Alexander and Dr. Kerry Dust of Cadham Provincial Laboratory. This study was funded/supported by the Public Health Agency of Canada and Genome Canada through the Canadian Public Health Laboratory Network COVID-19 Genomics Program (CCGP) and the Canadian COVID-19 Genome Network (CanCOGeN), respectively.

## Supporting information captions

**Figure 1:** Global nucleotide comparison of artificially generated genomes and their corresponding background clinical samples at a low sequencing depth. Heatmaps of the pairwise p-distance comparison of the delta background sequence (AY.25.1) contaminated with a delta contaminant sequence (AY.27). The different levels of contamination were shown for (A) medium (B) and high sequencing depths.

**Figure 2:** Global nucleotide comparison of artificially generated genomes and their corresponding background clinical samples at a low sequencing depth. Heatmaps of the pairwise p-distance comparison of an omicron background sequence (BA.1) contaminated with an alpha contaminant sequence (B.1.1.7). The different levels of contamination were shown for medium (A) and high (B) sequencing depths.

**Figure 3:** Phylogenetic tree and heatmaps showing single nucleotide variation at different positions of the SARS-CoV-2 genome for a delta variant (AY.25.1) contaminated with another delta variant (AY.27) sequence at contamination levels 1-10%, 20%, 30%, 40%, and 50% for (A) medium sequencing depth (25,000 reads) and (B) high sequencing depth sequence (50,000 reads).

**Figure 4:** Phylogenetic tree and heatmaps showing single nucleotide variation at different positions of the SARS-CoV-2 genome for an omicron variant (BA.1) contaminated with an alpha variant (B.1.1.7) sequence at contamination levels 1-10%, 20%, 30%, 40%, and 50% for (A) medium sequencing depth (25,000 reads) and (B) high sequencing depth sequence (50,000 reads).

**Figure 5.** Mutational profile comparison of SARS-CoV-2 genome for the clinical genomes to the artificially generated genomes for (A) MSD_SV and (B) (AY.25.1 contaminated with an AY.27 variant) sequence at contamination levels 1-10%, 20%, 30%, 40%, and 50%. (C) MSD_DV and (D) HSD_DV (BA.1 contaminated with a B.1.1.29 variant at contamination levels 1-10%, 20%, 30%, 40%, and 50%.

**Table 1A:** Quality control metrics comparison for artificially subsampled genomes of contamination by similar variants at a low sequencing depth – for all MSD_SV genomes.

**Table 1B:** Quality control metrics comparison for artificially subsampled genomes of contamination by similar variants at a low sequencing depth – for all HSD_SV genomes.

**Table 2A:** Quality control metrics comparison for artificially subsampled genomes of contamination by different variants at a low sequencing depth – for all MSD_DV genomes.

**Table 2B:** Quality control metrics for comparison of different SARS-CoV-2 sequences with a high number of reads (50,000 reads) at different levels of contamination.

## Notes

### Competing Interest Statement

The authors have declared no competing interest.

## References

1. Lupo V, Van Vlierberghe M, Vanderschuren H, Kerff F, Baurain D, Cornet L. Contamination in Reference Sequence Databases: Time for Divide-and-Rule Tactics. Front Microbiol. 2021;12. doi:10.3389/FMICB.2021.755101

2. Cornet L, Baurain D. Contamination detection in genomic data: more is not enough. Genome Biol. 2022;23. doi:10.1186/S13059-022-02619-9

3. Steinegger M, Salzberg SL. Terminating contamination: large-scale search identifies more than 2,000,000 contaminated entries in GenBank. Genome Biol. 2020;21: 115. doi:10.1186/S13059-020-02023-1

4. Park SY, Faraci G, Ward PM, Emerson JF, Lee HY. High-precision and cost-efficient sequencing for real-time COVID-19 surveillance. Sci Reports |. 2021;11: 13669. doi:10.1038/s41598-021-93145-4

5. Geoghegan JL, Douglas J, Ren X, Storey M, Hadfield J, Silander OK, et al. Use of Genomics to Track Coronavirus Disease Outbreaks, New Zealand. Emerg Infect Dis. 2021;27: 1317. doi:10.3201/EID2705.204579

6. Magalis BR, Ramirez-Mata A, Zhukova A, Mavian C, Marini S, Lemoine F, et al. Differing impacts of global and regional responses on SARS-CoV-2 transmission cluster dynamics. bioRxiv. 2020; 2020.11.06.370999. doi:10.1101/2020.11.06.370999

7. McLaughlin A, Montoya V, Miller RL, Mordecai GJ, Worobey M, Poon AFY, et al. Genomic epidemiology of the first two waves of SARS-CoV-2 in Canada. Elife. 2022;11. doi:10.7554/ELIFE.73896

8. Zhu Y, Liu M, Zhao W, Zhang J, Zhang X, Wang K, et al. Isolation of Virus from a SARS Patient and Genome-wide Analysis of Genetic Mutations Related to Pathogenesis and Epidemiology from 47 SARS-CoV Isolates. Virus Genes 2005 301. 2005;30: 93–102. doi:10.1007/S11262-004-4586-9

9. Yang Y, Peng F, Wang R, Guan K, Jiang T, Xu G, et al. The deadly coronaviruses: The 2003 SARS pandemic and the 2020 novel coronavirus epidemic in China. J Autoimmun. 2020;109: 102434. doi:10.1016/J.JAUT.2020.102434

10. Zhong NS, Zeng GQ. Our Strategies for Fighting Severe Acute Respiratory Syndrome (SARS). 101164/rccm200305-707OE. 2003;168: 7–9. doi:10.1164/RCCM.200305-707OE

11. Lu R, Zhao X, Li J, Niu P, Yang B, Wu H, et al. Genomic characterisation and epidemiology of 2019 novel coronavirus: implications for virus origins and receptor binding. www.thelancet.com. 2020;395: 565. doi:10.1016/S0140-6736(20)30251-8

12. Zhou H, Chen X, Hughes AC, Bi Y, Shi W. A Novel Bat Coronavirus Closely Related to SARS-CoV-2 Contains Natural Insertions at the S1/S2 Cleavage Site of the Spike Protein. Curr Biol. 2020;30: 2196–2203.e3. doi:10.1016/j.cub.2020.05.023

13. Wu F, Zhao S, Yu B, Chen YM, Wang W, Song ZG, et al. A new coronavirus associated with human respiratory disease in China. Nat 2020 5797798. 2020;579: 265 –269. doi:10.1038/s41586-020-2008-3

14. Zhu Z, Lian X, Su X, Wu W, Marraro GA, Zeng Y. From SARS and MERS to COVID-19: A brief summary and comparison of severe acute respiratory infections caused by three highly pathogenic human coronaviruses. Respir Res. 2020;21: 1 –14. doi:10.1186/S12931-020-01479-W/TABLES/4

15. Stoler N, Nekrutenko A. Sequencing error profiles of Illumina sequencing instruments. NAR genomics Bioinforma. 2021;3. doi:10.1093/NARGAB/LQAB019

16. Delahaye C, Nicolas J. Sequencing DNA with nanopores: Troubles and biases. PLoS One. 2021;16: e0257521. doi:10.1371/JOURNAL.PONE.0257521

17. Longo MS, O’Neill MJ, O’Neill RJ. Abundant Human DNA Contamination Identified in Non-Primate Genome Databases. PLoS One. 2011;6: e16410. doi:10.1371/JOURNAL.PONE.0016410

18. Breitwieser FP, Pertea M, Zimin A V., Salzberg SL. Human contamination in bacterial genomes has created thousands of spurious proteins. Genome Res. 2019;29: 954 –960. doi:10.1101/GR.245373.118/-/DC1

19. Lu J, Salzberg SL. Removing contaminants from databases of draft genomes. PLOS Comput Biol. 2018;14: e1006277. doi:10.1371/JOURNAL.PCBI.1006277

20. Goig GA, Blanco S, Garcia-Basteiro AL, Comas I. Contaminant DNA in bacteriARal sequencing experiments is a major source of false genetic variability. BMC Biol. 2020;18: 1–15. doi:10.1186/S12915-020-0748-Z/FIGURES/5

21. Bagheri H, Severin AJ, Rajan H. Detecting and correcting misclassified sequences in the large-scale public databases. Bioinformatics. 2020;36: 4699 –4705. doi:10.1093/BIOINFORMATICS/BTAA586

22. Malik P, Patel K, Pinto C, Jaiswal R, Tirupathi R, Pillai S, et al. Post-acute COVID-19 syndrome (PCS) and health-related quality of life (HRQoL)—A systematic review and meta-analysis. J Med Virol. 2022;94: 253–262. doi:10.1002/JMV.27309

23. Ramesh S, Govindarajulu M, Parise RS, Neel L, Shankar T, Patel S, et al. Emerging SARS-CoV-2 Variants: A Review of Its Mutations, Its Implications and Vaccine Efficacy. Vaccines. 2021;9. doi:10.3390/VACCINES9101195

24. David Nelson AR, Hazzouri KM, Lauersen KJ, Lomas MW, Amiri KM, Salehi-Ashtiani K. Large-scale genome sequencing reveals the driving forces of viruses in microalgal evolution. 2021 [cited 21 Jun 2023]. doi:10.1016/j.chom.2020.12.005

25. Maurya R, Shamim U, Chattopadhyay P, Mehta P, Mishra P, Devi P, et al. Human-host transcriptomic analysis reveals unique early innate immune responses in different sub-phenotypes of COVID-19. Clin Transl Med. 2022;12. doi:10.1002/CTM2.856

26. Boni MF, Lemey P, Jiang X, Lam TTY, Perry BW, Castoe TA, et al. Evolutionary origins of the SARS-CoV-2 sarbecovirus lineage responsible for the COVID-19 pandemic. Nat Microbiol 2020 511. 2020;5: 1408–1417. doi:10.1038/s41564-020-0771-4

27. Singh D, Yi S V. On the origin and evolution of SARS-CoV-2. Exp Mol Med 2021 534. 2021;53: 537–547. doi:10.1038/s12276-021-00604-z

28. Rahimi F, Talebi Bezmin Abadi A. Emergence of the Omicron SARS-CoV-2 subvariants during the COVID-19 pandemic. Int J Surg. 2022;108: 106994. doi:10.1016/J.IJSU.2022.106994

29. Markov P V, Ghafari M, Beer M, Lythgoe K, Simmonds P, Stilianakis NI, et al. The evolution of SARS-CoV-2. Nat Rev Microbiol |. 2023;21: 361–379. doi:10.1038/s41579-023-00878-2

